# Identifying causes of social evolution: The Price approach, contextual analysis, and multilevel selection

**DOI:** 10.1101/2020.08.14.233122

**Authors:** Christoph Thies, Richard A. Watson

## Abstract

Kin selection theory and multilevel selection theory are different approaches to explaining the evolution of social traits. The latter claims that it is useful to regard selection as a process that can occur on multiple levels of organisation such as the level of individuals and the level of groups. This is reflected in a decomposition of fitness into an individual component and a group component. However, the two major statistical tools to determine the coefficients of such a decomposition, the multilevel Price equation and contextual analysis, are inconsistent and may disagree on whether group selection is present. Here we show that the reason for the discrepancies is that underlying the multilevel Price equation and contextual analysis are two nonequivalent causal models for the generation of individual fitness effects (thus leaving different ‘remainders’ explained by group effects). While the multilevel Price equation assumes that the individual effect of a trait determines an individual’s relative success within a group, contextual analysis posits that the individual effect is context-independent. Since these different assumptions reflect claims about the causal structure of the system, the correct approach cannot be determined on general theoretical or statistical grounds but must be identified by experimental intervention. We outline interventions that reveal the underlying causal structure and thus facilitate choosing the appropriate approach. We note that the reductionist viewpoint of kin selection theory with its focus on the individual is immune to such inconsistency because it does not address causal structure with respect to levels of organisation. In contrast, our analysis of the two approaches to measuring group selection demonstrates that multilevel selection theory adds meaningful (falsifiable) causal structure to explain the sources of individual fitness and thereby constitutes a proper refinement of kin selection theory.

## 1 Introduction

When individual traits have effects on other individuals, individual fitness depends not only on self but also on the social environment, i.e., interaction partners. Kin selection theory (KS) deals with this problem by regarding the social environment as an external factor that, together with direct fitness effects of a trait, determines evolutionary dynamics with respect to selection. By assuming a certain correlation between trait value of an individual and average trait value of its social environment, e.g., through relatedness, Hamilton’s rule can be formulated and answers the question of whether a trait with direct and indirect effects increases or decreases in frequency given the organisation of the population, i.e., the parameter of relatedness *r* (Frank, 1997). In short, KS acknowledges indirect effects (for which it was developed) but focuses on how relatedness affects individual fitness and is indifferent to the levels on which selection acts.

Multilevel selection theory (MLS) differs from this picture in that it posits the social environment as a unit, e.g., the group, that can be subject to selection acting at a level above that of individuals (Wilson, 1975; Wade, 1976; Wade, 1978; Uyenoyama and Feldman, 1980; Wilson and Sober, 1989). The theory thus promotes the concept of a group from a mere collection of individuals targeted by similar selection pressures to a unit that has a causal role in the selection process. More precisely, MLS theory understands a group as a unit whose interaction with the selective environment - through properties of the group as a whole - causally affects the fitness of its individual subunits (Wade and Kalisz, 1990). This means that individual fitness is a composite quantity determined by two factors: the individual effect of the trait and an effect on the group that an individual is a part of, and via this group effect, on the individual itself. The MLS view is not in opposition with KS but merely highlights that selection at the group level may be part of a causal mechanism resulting in individual fitness differences and must be taken into account if we want to understand the source of individual fitness differences (Dugatkin and Reeve, 1994; Sober and Wilson, 1994). Put differently, while KS is content with determining inclusive fitness at the individual level, MLS claims that individual traits can have effects that are best understood as group effects. Note that KS and MLS make the same predictions about trait-frequency dynamics on the individual level because selection at higher levels entails selection at lower levels and KS interprets all selection in individual terms. The explanatory goals of KS itself (Okasha, 2015; Marshall, 2016) derive largely from the goal of establishing inclusive fitness as a quantity maximised by evolution (Hamilton, 1964). Here, we refer to KS as a model that is free of assumptions regarding the level of selection in the sense that KS subsumes all selection at the individual level while MLS deviates from this model by assigning selection to several levels. While MLS aims to analyse the proximate causal structure of selection at multiple levels of organisation, KS establishes the direction of trait-frequency change based on individual fitness consequences of the trait and the relatedness structure of the population.

The distinction between individual effects and group effects of individual traits presents MLS with a problem not encountered by KS: how can the presence of a group effect be detected empirically/statistically and how can the strength of the group effect be quantified in comparison to the individual effect of the trait. After all, the claim that group effects determine individual fitness can only be of use if such effects can be detected empirically. To give an example, Eldakar et al. (2010) claim that the fitness of male water striders *Aquarius remigis* organised into patches (also referred to as social environments or groups) depends on two components that are both affected by an aggressiveness trait individually expressed by the males. The individual component is given by the positive effect of aggressiveness on fitness mediated by mating success which is higher for more aggressive males that secure more mating opportunities than less aggressive males (Sih and Watters, 2005). The group component of individual fitness, on the other hand, arises from a different causal pathway and represents a negative effect of aggressiveness on fitness. Since the harassment experienced by females on a patch reflects the cumulative male aggression level on that patch and females tend to avoid harassment by escaping their current patch, the trait has a negative effect on patch productivity by decreasing the number of females on the patch and therefore the reproductive resources of all males on that patch. If such a decomposition into causes of individual fitness is to be useful, this decomposition must be empirically accessible in the sense that fitness is quantitatively given as a function of an individual component and a group component. This is possible only with a valid method of measuring the decomposition in empirical data.

Two methods for carrying out a quantitative decomposition of individual fitness into an individual component and a group component have received particular attention in the literature (Heisler and Damuth, 1987; Goodnight, Schwartz, and Stevens, 1992; Frank, 1998; Okasha, 2006; Sober 2011, McLoone, 2015): the multilevel Price equation and contextual analysis which, following Okasha, we refer to as the ‘Price approach’ and the ‘contextual approach’, resp. However, the partitions of individual fitness given by the two methods are different in general. In particular, there are cases in which the multilevel Price equation claims the absence of group effects while contextual analysis claims their presence and vice versa.

The inconsistency between the two approaches is problematic because proponents of MLS argue that the distinction between individual effects and group effects is not just a statistical exercise but reflects a separation of causal pathways in the biological system under study as described above. While one causal pathway emanating from the individual trait is proposed to affect only individual aspects of fitness (the fitnesses of the bearer and its interaction partners), a different pathway is claimed to relate the trait with properties of the group as a whole and hence with a group component of individual fitness. Since the desired decomposition must reflect the underlying biological reality, two methods of decomposition that yield different answers cannot both be correct (Sober, 2011). Previous attempts at resolving these discrepancies have been inconclusive, leaving theorists and empiricists applying multilevel selection theory in the unfortunate situation that, even among proponents of multilevel selection theory, there is no unanimously agreed upon method for measuring the strength of group selection in the simplest additive cases (Eldakar et al., 2010; Clarke, 2016). Given that multilevel selection is still contentious in traditional evolutionary theory, and even the proponents of MLS have been unable to agree on a measure (and thus even unable to agree whether a group effect is present or not in a particular case), this may suggest that MLS is not well understood and should be abandoned in favour of kin selection theory.

The aim of this paper is to show that the essential difference between the Price approach and contextual analysis lies in the causal structure each method posits as underlying the observed measurements of individual fitness. Briefly, while contextual analysis assumes that the individual component is determined by direct fitness effects of the trait only, the Price approach sees the individual component as a result of within-group competition and duly assumes it to be affected by the trait values of group mates (indirect fitness effects). Put differently, contextual analysis assumes that the individual effect of the trait is absolute in the sense that it is independent of the social environment. The Price approach, on the other hand, assumes that the trait affects the competitiveness of its bearer so that its fitness effect is relative in the sense that it depends on the social environment (see cartoon example in Figure 1). This difference leads to different remainders to be explained by group effects and thereby to different measurements of the strength of group selection. Recognising that the difference between the two approaches arises from a difference in the underlying model of reality enables us to see how to determine which of the two approaches is correct in a given case, i.e., the one whose underlying model reflects the causal structure of the system that is being studied. In particular, the applicability of the two approaches depends on the biological scenario at hand and cannot be made on theoretical grounds.

**Figure 1:**
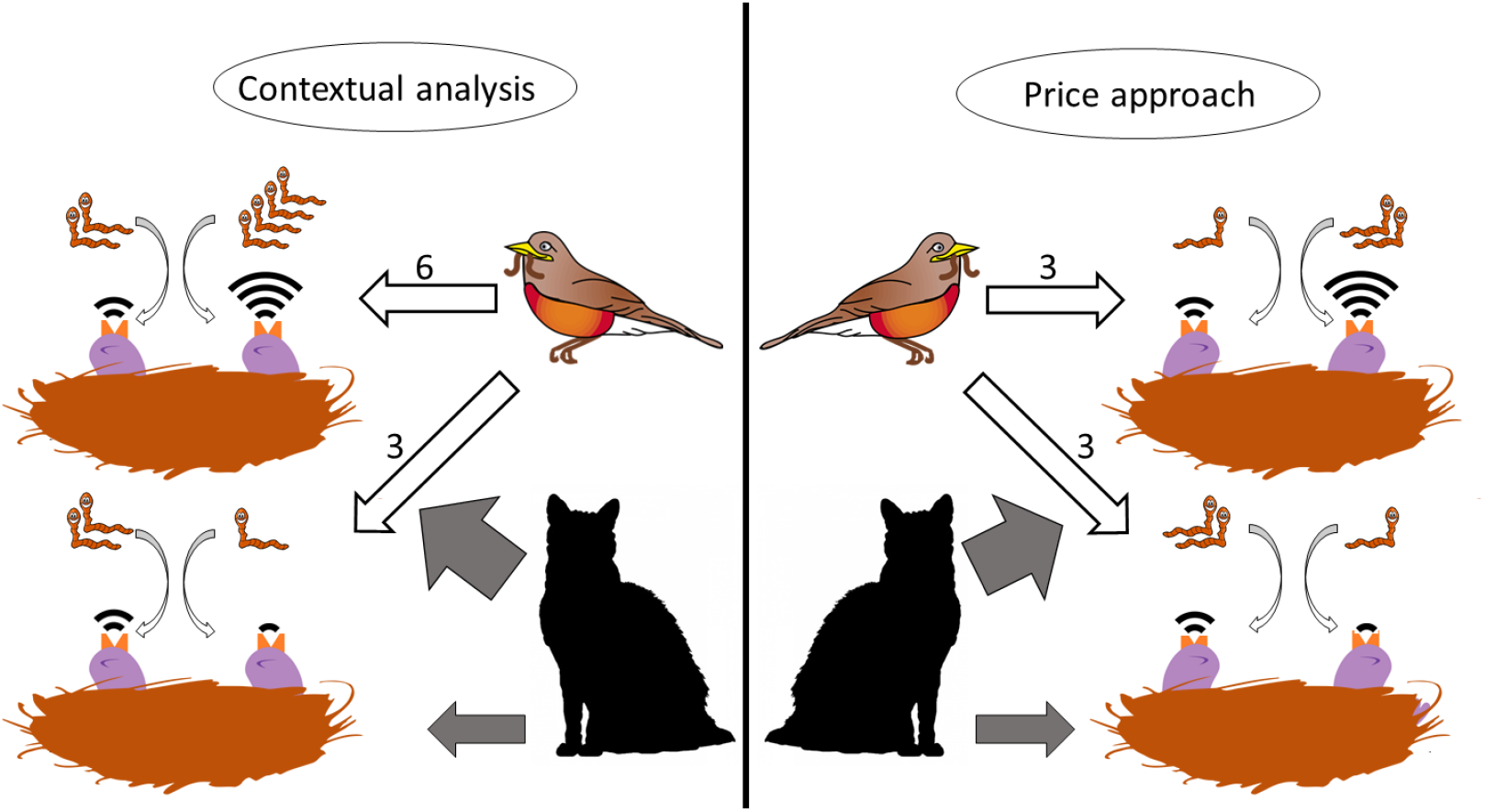
The different underlying assumptions of the Price approach and contextual analysis. A chick’s fitness depends on two factors: the number of worms it receives (individual component) and the risk of its nest being discovered by a cat (group component). The trait ‘begging volume’ has effects on both fitness components. The louder a chick begs the more worms it receives and the more likely it is that its nest is found by a cat. But how much group effect is there exactly? This depends on the underlying assumptions of how the trait affects the individual component. Contextual analysis assumes an individual component that is independent of the nest mates so that a chick gets a number of worms determined by its own begging volume only. This means the total begging in a nest affects predation risk but also affects the total worms received by that nest (the louder upper nest receives 6 worms, the lower more quiet nest only 3). The Price approach assumes an individual component that results from competition within the nest so that a chick gets a number of worms determined by the ratio of its begging volume to the total begging volume in the nest. This means that the total begging in a nest affects predation risk as before but does not affect the total worms received by that nest (both nests receive 3 worms).

This paper is organised as follows. First, we show that both contextual analysis and the Price approach can be interpreted in terms of causal graphs that describe how each of the approaches models the dependence of individual and group component of fitness on individual and group trait. We then compare the two approaches using these causal graphs. This allows us to illustrate very clearly why the two approaches give different answers with respect to the strength of group selection. In addition to identifying the source of the discrepancy, our analysis identifies an experimental intervention that reveals which, if any, of the two approaches is correct and shows that neither is always correct. Indeed, the correct approach depends on causal mechanisms in the biological system that cannot be determined based on the distribution of individual fitness over individual trait and group trait without experimental intervention.

We see the wider relevance of this work in two points. First, we suggest a resolution to the problem of inconsistency between the Price approach and contextual analysis. We present, to our knowledge for the first time, a complete and unifying description of the difference between the two approaches, including a method for choosing which method to use in an empirical scenario.

Settling this issue helps strengthen the theoretical core of multilevel selection. Second, our analysis demonstrates how the application of MLS to a biological scenario requires and formalises an understanding of the system that is not implied by KS. More precisely, MLS introduces a layer in the causal structure that cannot be deduced from the reduced theory. Identifying this causal substructure requires an intuitive inspection of the empirical system. We thus demonstrate transparently, and in the simplest case of additive interactions, how MLS represents a non-reductionist refinement of KS with respect to the causal structure of selection.

## 2 Model

### 2.1 Fitness, selection, and the Price equation

The evolutionary model in which we frame our arguments is as simple as possible whilst being able to support the features we set out to discuss. Individuals are defined by their allele at a biallelic locus, with the two alleles representing the presence and absence of a trait, which also defines their phenotype (denoted by *x* with *x* = 0 if the trait is absent and *x* = 1 if the trait is present) and replicate asexually without mutation. A population of individuals is partitioned into non-overlapping groups of equal size such that an individual interacts equally with all members of its group (the assumptions on group size and disjointness are made for convenience only). The absolute fitness of an individual determines per capita growth rate and is a function of its own trait as well as the group trait, i.e., the mean trait value of the individuals in a group, but not a function of other properties of the population (absence of, e.g., global frequency dependence). Taking a causalist stance, we assume that this function is deterministic rather than a statistical abstraction from data (Otsuka, 2016) and stable in its functional form (i.e., the selective environment that determines fitness in interaction with the phenotype is not changing (Wade and Karlisz, 1990)). Moreover, we assume that the fitness function is additive such that

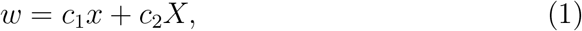

where *w* denotes the fitness of an individual with phenotype *x* and group phenotype *X*, and coefficients 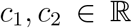 (Okasha, 2006). This notation corresponds to the method of direct fitness or neighbour-modulated fitness in KS (Taylor, Wild, and Gardner, 2007). *c*_1_*x* represents the direct fitness effect of the trait on its bearer, *c*_2_*X* the indirect fitness effect on trait bearers’ interaction partners.

The process of selection in a population is given by the change in trait frequency according to the Price equation without mutation

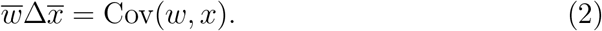

Note that we do not assume that groups themselves replicate or can be assigned group fitness over and above the fitness of the individuals that constitute a group. Our model is therefore of MLS1 type in the sense of Heisler and Damuth (1987), i.e., the focus of the analysis is on individuals, group trait and group fitness are averages of the corresponding quantities of the individuals within the group.

The starting point for the analysis of selection in a population in terms of MLS is the observation that an aspect of selection acts on groups as a whole. This means that individual selection is in part determined by the group trait *X* because selection favours groups with high (or low) group trait. In particular, this aspect of selection is the same for all members of a group and is captured by the process by which some groups contribute more offspring to the next generation than others due to differential proliferation and extinction (Uyenoyama and Feldman, 1980; Wade and Goodnight, 1998). Note that it makes no difference to the change in trait frequency whether (an aspect of) selection acts on the group as a whole or on all group members individually but in the same way. However, the aim of MLS is not only the prediction of outcomes but also the attainment of a causal understanding of the selection process (Sober and Wilson, 1994). The observation that selection depends on a property of the group as a whole is reflected in a decomposition of individual fitness into a component that is common to all group members because they share the same group property and a component that differs among group members (Sober, 1980; p.107). The trait that - directly for its bearer and indirectly for its bearer’s group mates - determines individual fitness therefore has an individual effect and a group effect

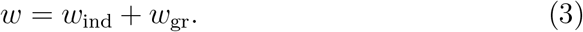

A few remarks concerning Equation (3) are in order. The purpose of this decomposition is to explicitly and formally acknowledge the basic tenet of MLS that fitness (here at the individual level) is determined not only by how the individual interacts with its selective environment but also by how the individual’s group interacts with the selective environment on a level above that of the individual. We introduce *w_gr_* to formally capture fitness effects that result from the interaction of the group as a whole with the selective environment. The quantities *w_ind_* and *w_gr_* are proxies for the effects of causal processes, the former for processes that affect individual fitness specifically for each individual, the latter for processes that affect the group. In the example pictured in Figure 1, fitness of an individual chick is determined via two differing causal pathways whose separation is expressed mathematically by Equation (3): *w_ind_* is the aspect of fitness that is determined by how well-fed the chick is, while *w_gr_* is the aspect of fitness that is determined by how likely the chick is to be found by a predator (together with its nest mates). To apply this decomposition empirically it may be possible to identify measurable proxies for the components. In this example, this might be the amount of food a chick receives over some period of time for *w_ind_* and the number of attempted raids on the nest for *w_gr_*. It should be noted that the decomposition in Equation (3) is additive for simplicity only. While an MLS analysis always implies a decomposition of fitness into contributions from various levels, this decomposition is, generally, not additive. The formalisation in terms of causal graphs introduced below does not require additivity.

Since the Price equation is linear in the fitness argument, the decomposition expressed in Equation (3) corresponds to a decomposition of the strength of selection itself

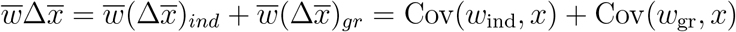

In order to make quantitative statements about the strengths of group selection vs. individual selection, an MLS analysis must determine the components in this decomposition. However, while individual trait and fitness as well as aggregates thereof can be measured directly, individual effect and group effect, or their covariance with the individual trait, are generally not amenable to direct measurement. The multilevel Price equation and contextual analysis are two methods of obtaining *w*_ind_ and *w*_gr_ by statistical means given individual traits and fitnesses (Okasha, 2006).

### 2.2 Contextual analysis and the Price approach

Equation (1) partitions individual fitness into the effects of the individual trait and the group trait. It describes how two phenotypic traits combine to yield another trait of the individual, namely absolute fitness. Contextual analysis (Heisler and Damuth 1987; Okasha, 2006; note that here and the following we refer by ‘contextual analysis’ to the standard regression on the untransformed variables *x* and *X*, as is customary in discussions on the issues reported here) takes effects of the group trait in Equation (1) as indicating group selection. Strictly speaking, *c*_2_ ≠ 0 in Equation (1) implies the potential of the trait to undergo group selection conditional on the existence of group-trait variation between groups (Wolf et al., 1999; see McLoone (2015) for a discussion of this difference). We regard group effects on fitness as more fundamental than a concept of group selection itself as the former do not depend on properties of a population but reflect causal processes that increase or decrease reproductive success of an individual situated in a group context vis-à-vis a specific selection regime that in turn determines individual fitness. Group effects can lead to group selection if, in a specific population, they generate fitness differences between individuals. This requires Var(*X*) ≠ 0, for if Var(*X*) = 0 all individuals have the same group trait and are therefore subject to the same group effects. The Price equation maps a fitness function understood as superposition of fitness effects of the variables that causally determine fitness to selection in a population in which individual fitness is given by the assumed fitness function (Figure 2).

**Figure 2:**
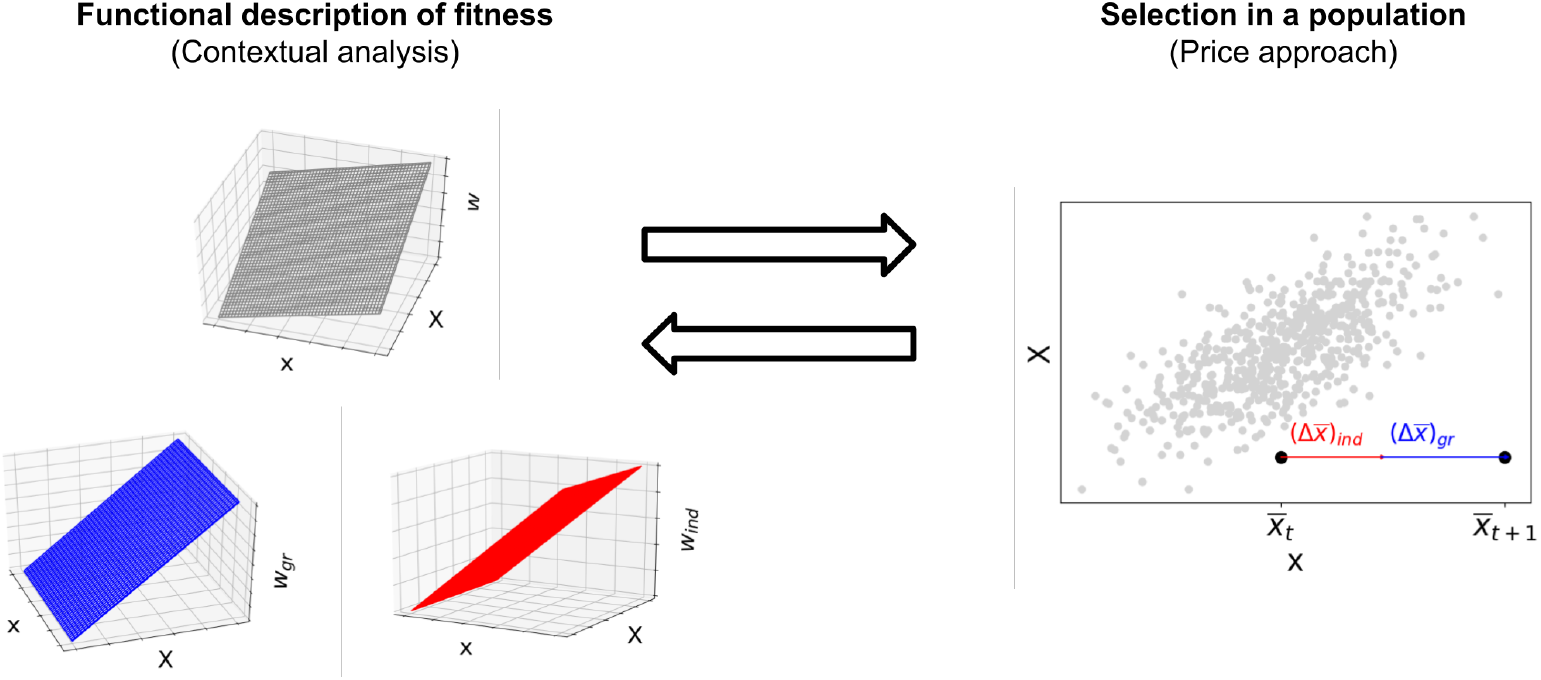
Contextual analysis decomposes fitness into an individual and a group component while the multilevel Price equation decomposes selection into individual and group selection. The standard Price equation without transmission bias maps a functional description of fitness to a process of selection in a population of individuals whose fitness is defined by this functional description. Since the Price equation is linear in the fitness function, a decomposition of fitness into the summands individual and group effects yields a corresponding and unique decomposition of selection into the summands individual and group selection.

The Price approach to multilevel selection (Price, 1972; Okasha, 2006) rests on the partition of selection itself given by the multilevel expansion of the Price equation (2)

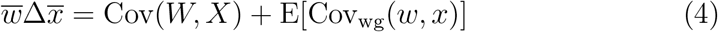

and posits that a population is undergoing group selection if the first term in Equation (4) is non-zero. In light of our remarks regarding group effects and group selection above, the Price approach and contextual analysis therefore decompose different quantities and are not directly comparable. However, this difference is superficial as the partition of fitness effects given by contextual analysis corresponds to a partition of selection and the partition of selection given by the multilevel Price equation corresponds to a partition of fitness effects. Contextual analysis, i.e., the functional representation of fitness in Equation (1), determines selection according to Equation (2) for a population that is partitioned into groups: given a population of individuals *i* ∈ 1,…, *n* with fitnesses

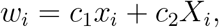

where *X_i_* is the trait of the group the *i*^th^ individual is part of, the change in mean trait value in the population follows from Equation (2) as (Okasha, 2004)

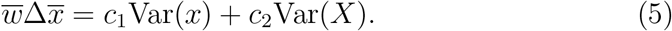

Thus the decomposition of fitness into individual and group effects given by contextual analysis corresponds to a decomposition of selection whose components, according to contextual analysis, represent the component of individual selection *c*_1_ Var(*x*) and the component of group selection *c*_2_Var(*X*).

Conversely, the components of individual selection and group selection according to the Price approach for a population with non-vanishing variance within and between groups correspond to a decomposition of individual fitness into a component of individual effects and group effects. To see how, note that with *w* = *c*_1_*x* + *c*_2_*X*,

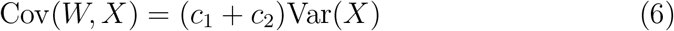

(Okasha, 2006; p.89). Using Equation (5) and Equation (6), the decomposition according to Equation (4) is

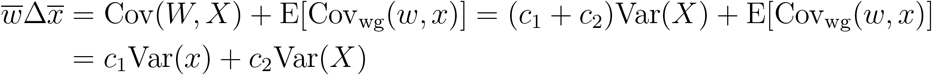

and therefore

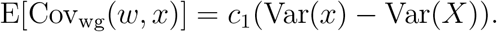

Hence the decomposition of fitness

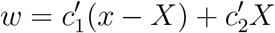

corresponds to the decomposition of selection

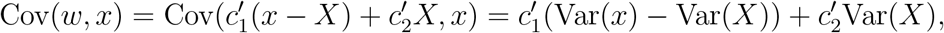

where 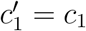 and 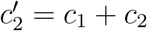.

Through this correspondence both contextual analysis and the Price approach yield decompositions of fitness effects as well as of selection (see Table 1). Note that the possibility of conducting contextual analysis with respect to the variables *x* – *X* and *X* rather than *x* and *X* – the former choice of variables being equivalent to the Price approach, the latter to contextual analysis – is discussed in Heisler and Damuth (1987) along with examples of circumstances under which this might be causally adequate.

**Table 1:** The decompositions of contextual analysis and the Price approach as individual and group fitness effects *w* = *w*_ind_ + *w*_gr_ and as components of selection. The parameters of contextual analysis and the Price approach are linked by the equations 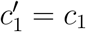 and 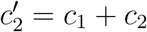.

|  | Fitness | Selection |
| --- | --- | --- |
| | Individual component + group component<br>$w = w_{\text{ind}} + w_{\text{gr}}$ | Individual selection + group selection<br>$\Delta\bar{x} = (\Delta\bar{x})_{\text{ind}} + (\Delta\bar{x})_{\text{gr}}$ |
| Contextual analysis | $c_1x + c_2X$ | $c_1\text{Var}(x) + c_2\text{Var}(X)$ |
| Price approach | $c'_1(x - X) + c'_2X$ | $c'_1(\text{Var}(x) - \text{Var}(X)) + c'_2\text{Var}(X)$ |

### 2.3 Causal intuitions underlying an MLS analysis

A core idea of social evolution is that an individual trait of social organisms has fitness effects not only on its bearer but also on the social environment of the bearer. Common to paradigmatic examples of group selection is an individual trait with effects that change individual fitness homogeneously across the group such that these effects are best viewed as group effects (Sober, 1980). For the water striders described in Eldakar et al. (2010) the exodus of females from patches with high levels of aggressiveness is a group effect of the trait ‘aggressiveness’ in males. This group effect is negative because group productivity is assumed to decrease with the number of females on a patch as females provide reproductive resources. The causal interpretation of the trait refers to proximate fitness effects of the trait and involves the individual as well as the group it is in but not other groups or the population as a whole. Therefore the causal interpretation takes place on the fitness side rather than on the selection side in Figure 2.

Since we assume that fitness is an effect of the individual/group trait an individual exhibits, we can read the equations in the left column of Table 1 as structural equations that determine fitness given the traits. By the assumption on the additivity of interactions these equations are linear. The interpretation of structural equations is aided by the use of causal graphs, more precisely, directed acyclic graphs with causal rather than correlational interpretation (Pearl, 2009). Figure 3 shows the causal graphs corresponding to the structural equations in Table 1. Since the components *w*_ind_ and *w*_gr_ reflect quantities that refer to processes occurring in the biological system studied, the causal graphs constitute models of the underlying reality. For example, the group effect of the aggressiveness trait in water striders is given by the propensity of females to remain on the focal patch and this propensity is a function of mean male aggressiveness in the patch (this function is linear by assumption), i.e., the group trait *X*. The non-equivalence of the causal graphs (a) and (b) in Figure 3 reflects a difference in how the individual/group components of individual fitness depend on individual/group trait. It should be noted that the factors *x* and *X* are not strictly independent as suggested by omitted arrows between *x* and *X* in Figure 3. Since the group phenotype is generated collectively by all individuals within a group, *x* does affect *X*. The arrows are omitted in Figure 3 because our arguments focus on that part of the causal structure that determines fitness. Details of how the interaction of individual phenotypes gives rise to the group phenotype are not relevant for the present discussion.

**Figure 3:**
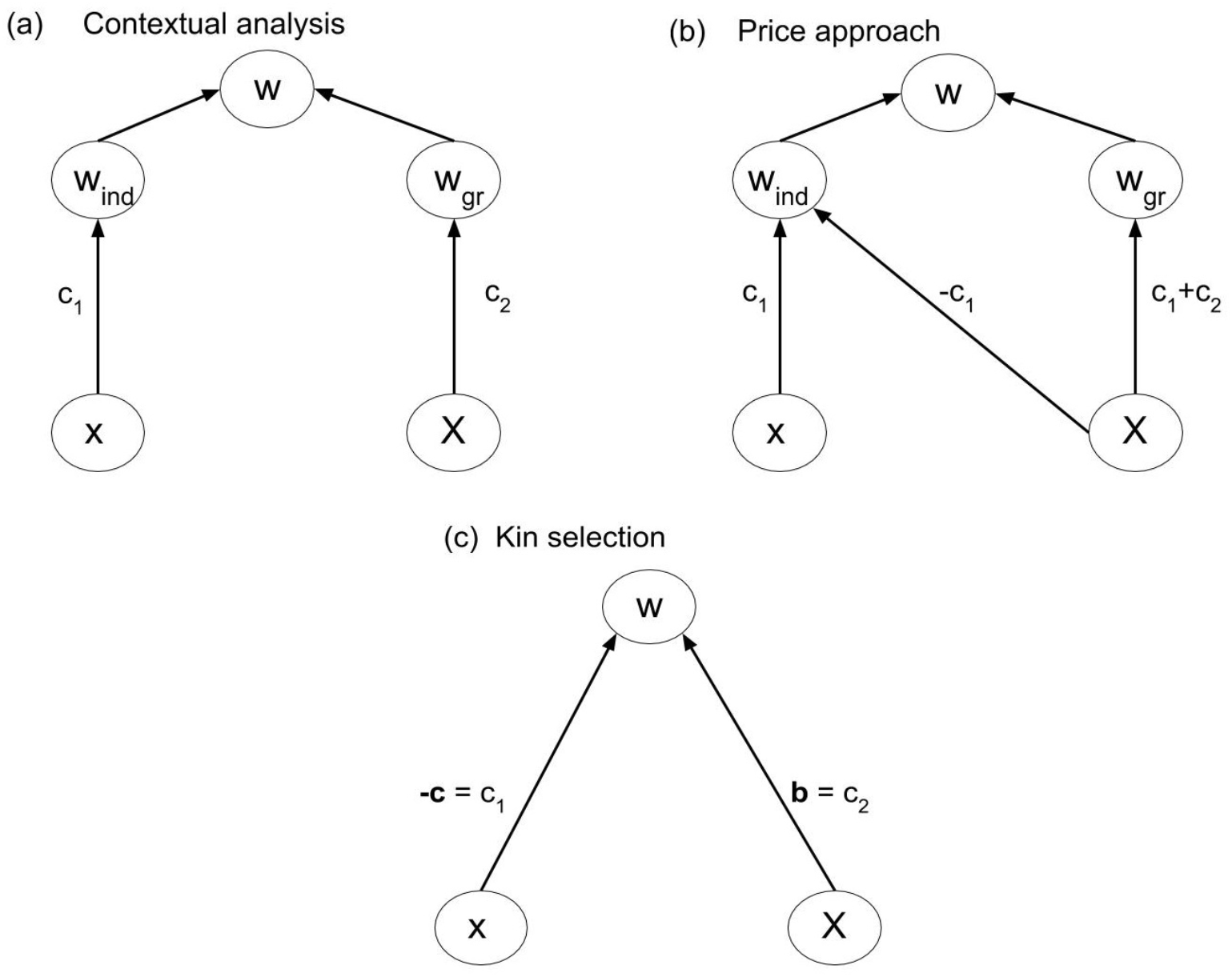
Causal graphs showing the interdependence of the variables *x* (individual trait), *X* (group trait, i.e., group mean of individual trait), *w* (individual fitness), *w*_ind_ (individual component of individual fitness), and *w*_gr_ (group component of individual fitness) (a) Contextual analysis assumes an absolute individual effect of the trait. (b) In the Price approach, the trait is assumed to have a relative effect in the sense that the trait affects fitness depending on the trait expression of other members of the group. (c) In contrast, kin selection theory acknowledges the possibility of indirect effects in addition to direct effects but makes no further assumptions on the causal structure. In KS, it is customary to denote the direct effect of the trait on its bearer by – *c* and the indirect effect by *b*. The parameter of relatedness *r* represents the correlation between *x* and *X* and is not pictured in the graph because we focus on selection rather than on properties of group composition. Also the effect of individual phenotype on group phenotype has been omitted, see text.

The model of fitness underlying contextual analysis (panel (a) in Figure 3) is based on the assumption that the individual component and the group component of fitness are determined only by the individual trait and the group trait, resp. This means that fitness differences within groups, i.e., differences in the individual component, are due to the individual trait and independent of the group trait. In that sense contextual analysis assumes the individual effects of the trait to be absolute, i.e., independent of group context. In contrast, the Price approach assumes that the group trait also affects the individual component of fitness in a specific way (see the path coefficients in Figure 3). This effect of the group trait on the individual component is equivalent to the assumption that fitness differences within groups are due to competition between group members in which the individual trait determines competitiveness of an individual. Indeed, the functional representation of fitness according to the Price approach from Table 1

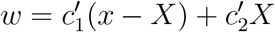

shows that the individual component sums to zero over each group and that individuals with higher-than-average trait have a positive individual component (negative if 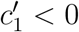). In other words, the trait affects individual fitness not by generally increasing or decreasing its bearer’s fitness but by increasing or decreasing its bearer’s competitive ability within the group. We discuss examples of these differences in the next section.

## 3 Results

### 3.1 Cases of disagreement

When comparing the Price approach and contextual analysis it should be kept in mind that both aim to quantify group selection and therefore start with the intuitive identification of an effect of the trait on a group-level property that affects fitness of all individuals within a group homogeneously. In the water strider example, an MLS analysis is based on the assertion, intuitively acquired by inspection of the empirical system, that the group mean of the trait ‘aggressiveness’ affects the number of females in a group and therefore the productivity of the group as a whole. This assertion is independent of the subsequent choice of statistical approach to quantifying the strength of group selection. Contextual analysis and the Price approach therefore agree on the nature of the group effect on fitness *w*_gr_ and on the mechanism bringing forth this effect, though not on its magnitude. The difference between the two approaches lies in the question of which factors affect the individual component of fitness, i.e., which factors are responsible for within-group differences in fitness.

The problem cases for contextual analysis and the Price approach discussed by Okasha (2006) and others (Heisler and Damuth, 1987; Sober, 2011; Goodnight, 2015) reveal issues with the two approaches because the intuition about the level on which fitness effects occur is inconsistent with the verdict of one of the approaches with respect to the strengths of individual and group selection. This intuition is best understood in terms of fitness effects and not in terms of selection because it is based on a mechanism that mediates an effect of the trait on the group component of absolute fitness and is therefore independent of composition and organisation of the population as a whole. Changing a patch of water striders to exhibit a lower level of the group trait ‘aggressiveness’ increases group fitness because less females will flee the patch. This causal explanation of the biological scenario is the core of an MLS analysis and it is independent of other patches and selection dynamics in the overall population. We conclude that the intuition with regard to the levels on which selection acts is about the mechanisms and not about frequency changes in the population. Accordingly, the following discussion is couched in terms of the left-hand side of Figure 2.

In the following examples, we determine the coefficients *c*_1_, *c*_2_ of the kin selection model (Figure 3 (c)) and discuss their interpretation in terms of the refined models provided by contextual analysis and the Price approach (models (a) and (b) in Figure 3).

#### 3.1.1 Non-social trait

A trait is non-social if the fitness of an individual does not depend on the trait values of its interaction partners (group mates) (Okasha, 2006) so that *c*_2_ = 0 and *w* = *c*_1_*x* (*c*_1_ ≠ 0 unless the trait is altogether neutral) in Figure 3. Intuitively, a trait of this type cannot be subject to group selection, because it has no fitness effects on its bearer’s interaction partners and therefore cannot affect the group component of fitness. However, the causal graph that represents the assumptions of the Price approach (Figure 3 (b)) shows an effect of group trait on group component of fitness with weight *c*_1_ + *c*_2_ = *c*_1_ and therefore detects group selection where intuitively there is none. Group effects of this type have been called cross-level by-products (Okasha, 2006) and will be discussed in a later section. Note that the causal graph underlying contextual analysis correctly shows the absence of group effects.

#### 3.1.2 Soft selection

The tension between the Price approach and contextual analysis is reversed in the case of soft selection (Wade, 1985; Goodnight, Schwartz, and Stevens, 1992; Débarre and Gandon, 2011). Briefly, soft selection occurs in a group-structured population if mean individual fitness is homogeneous across groups, i.e., if all groups have the same reproductive output. Soft selection models situations in which individuals of each group share a fixed resource and the trait under soft selection determines how an individual fares in the within-group competition for this resource. The group trait determines competitiveness of the group, i.e., mean competitiveness of its members, in the sense that an individual has lower fitness in a competitive group than in a group with low group trait. Soft selection is intuitively considered to be free of group selection (Wade, 1985; Okasha, 2006; Sober, 2011). The trait has no effect on the group level because changing the trait value of an individual in a group has no homogeneous fitness effect within the group as the change has no consequences for mean group fitness but merely changes the outcome of the within-group competition. It is easy to see that a kin selection model of soft selection takes the form *w* = *c*_1_*x* – *c*_1_*X*, i.e., *c*_2_ = −*c*_1_, with *c*_1_*x* > 0 (resp. *c*_1_ < 0) if a higher trait value implies higher (resp. lower) competitiveness.

The interpretation of these parameters according to the Price approach yields that the edge from *X* to *w*_gr_ has weight *c*_1_ + *c*_2_ = 0 in the causal graph (b) in Figure 3. The Price approach correctly detects the absence of group selection in this example. However, contextual analysis mistakes the effect of the group trait on fitness as an effect on the group component of fitness according to the causal graph (a) in Figure 3.

Though most researchers that engaged with the problem of inconsistency between contextual analysis and the Price approach seem to agree that no group selection occurs in soft selection, some have argued to the contrary. Goodnight, Schwartz, and Stevens (1992) regard soft selection as an example of group selection since an individual’s fitness depends on the trait of the group of which it is a member. We agree that individual fitness depends on the group trait but this effect of the group trait on fitness is an individual effect (the diagonal arrow in Figure 3 (b) targets *w*_ind_) that represents within-group competition. In soft selection, there is no group effect since the trait does not influence group fitness.

#### 3.1.3 Genotypic selection with meiotic drive

Okasha (2004) introduces ‘frameshifting’ as a desirable property of a general theory of multilevel selection. The theory is capable of frameshifting if it formalises features of group selection in such a way that they hold by analogy whenever the hierarchy given by the group/individual relation is instantiated at other levels of organisation. The treatment of genotypic selection with meiotic drive in MLS terms is relevant in that context because it tests the ability of MLS to frameshift to levels below the level of organisms. Following Wilson (1990), Okasha (2004) discusses diploid population genetics as an example of multilevel selection where alleles correspond to individuals and diploid genotypes to groups. In this analogy, group effects on allelic fitness are due to genotypic fitness, i.e., organismic fitness of the organism with a specific genotype, and individual effects are due to meiotic drive that creates within-group fitness differences between alleles.

Given the intuition that individual selection as well as group selection is at work in genotypic selection with meiotic drive, the expectation with respect to a decomposition of fitness into individual and group effects is clearly that group selection must always be present whereas individual selection is brought about by unfair meiosis. However, it is easy to see using specific fitness functions that contextual analysis doesn’t agree with intuition in this case. In particular, for a fitness function that depends on individual trait and group trait in such a way that allelic fitness is independent of the genetic background despite unfair meiosis and dependence of fitness on the group trait (*c*_1_ ≠ 0, *c*_2_ = 0), contextual analysis concludes the absence of group selection. The Price approach, in contrast, reaches the correct conclusion that individual fitness is given by 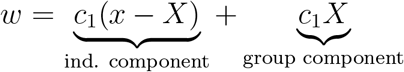 and therefore that both components of selection are non-zero.

Okasha’s conclusion that the “covariance approach [i.e., the Price approach] appears to frameshift down quite well, the contextual approach very badly” (Okasha, 2004; p.498) is thus readily explained by the viewpoint developed so far: unfair meiosis corresponds to the zero-sum game of within-group competition. This is precisely the causal structure assumed by the Price approach.

### 3.2 Cross-level by-products

A core assumption of MLS theory is that a trait an individual expresses may affect properties of its group as a whole and therefore group fitness (i.e., mean individual fitness in a group). This effect is captured by the group component *w*_gr_ of individual fitness. However, group fitness, in MLS1, is the average individual fitness in a group and therefore comprises not only the group component but also the average individual component *w*_ind_. This is problematic because the part of group fitness that entails selection on the group property caused by the trait is *w*_gr_ only. The contribution of *w*_ind_ to group fitness is called a cross-level by-product (Okasha, 2006) because it represents fitness of the individuals that constitute the group, i.e., the lower level, rather than fitness that is a property of the group as a whole, i.e., the higher level. Intuitively, a group with many individually fit members seems more fit than a group with few individually fit members even when the group component *w*_gr_ and therefore the fitness vis-à-vis group selection that is to be quantified is the same for both groups. The non-social trait case discussed above is a good example of this effect. Since there is no group property for group selection to act on in this case, group fitness comprises solely of individual fitness from the level below and therefore consists entirely of cross-level by-products.

To see how contextual analysis and the Price approach handle cross-level by-products assume that individual fitness is given by the expression *w* = *c*_1_*x*+*c*_2_*X*. The decomposition of group fitness *W* = (*c*_1_ + *c*_2_)*X* into a component due to group effects and a component due to individual effects depends on the causal structure and therefore differs between the two approaches. While contextual analysis partitions group fitness into individual and group component as 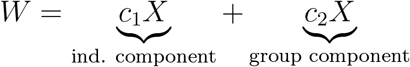 the decomposition according to the Price approach yields only a group component, *W* = (*c*_1_ + *c*_2_)*X*. For a non-social trait (*c*_1_ ≠ 0, *c*_2_ = 0) the Price approach mistakenly traces group fitness entirely back to a non-existing group effect, whereas contextual analysis correctly assigns group fitness to the individual effect. The fact that contextual analysis handles cross-level by-products correctly in the non-social trait case has led Okasha to conclude that contextual analysis is “on balance preferable” (Okasha, 2006; p.99) to the Price approach. However, it should be noted that in the soft selection case (*c*_1_ = −*c*_2_) contextual analysis decomposes group fitness as

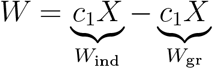

and hence detects cross-level by-products of magnitude *c*_1_*X* even though cross-level by-products are absent since the individual components of fitness sum to zero in each group.

In their study on multilevel selection in water striders, Eldakar et al. (2010) choose contextual analysis for quantifying group selection because contextual analysis controls for “potential cross-level byproducts” (Eldakar et al., 2010; p.3186). However, as we have seen, contextual analysis does not correctly account for cross-level by-products automatically. Which of the two approaches is correct depends on the kind of individual selection that acts on the system, i.e., the causal structure underlying fitness. In this case, the causal graphs (a) and (b) in Figure 3 both seem possible. Recall that aggressiveness in male water striders is hypothesised to have an effect on the individual component of fitness (aggressive males secure more mating opportunities than non-aggressive males) and on the group component of fitness (patches with higher aggression levels have fewer females). Contextual analysis assumes that the individual component is independent of the group trait: in addition to the group component shared by all males in a group each male has an individual component that is determined by its trait and independent of the group trait. Another, perhaps more plausible, assumption underlies the Price approach: the group trait determines the number of females on a patch and this reproductive resource is distributed to the males according to their competitiveness. We will discuss an experimental intervention that would reveal the correct underlying causal structure in the next section.

### 3.3 Determining the preferable approach

Several authors have discussed the question which of the two approaches is preferable in general (Okasha, 2006; Sober, 2011; McLoone, 2015). However, even the most extensive discussion of this question (Okasha, 2006) has been inconclusive in the sense that in light of the problematic cases discussed above neither can be endorsed unreservedly. We argued that a general preference cannot be justified as the essential difference between the two approaches lies in non-equivalent assumptions about the causal structure of the biological system which, as the problematic cases demonstrate, may be either of the two. However, our reduction of the difference between the Price approach and contextual analysis to a difference between their respective causal graphs has the benefit that experimental interventions that reveal the correct causal structure and with it the correct approach can easily be derived from the causal graphs (Pearl, 2009). Note that while we argue that the suggested interventions in principle reveal the correct structure we do not claim that such interventions are feasible for a given biological system. Moreover, while the two approaches discussed here are the main approaches to measuring the strength of group selection, it may well be possible that neither is suitable in a given scenario. We will discuss this and other limitations of this work below.

Imagine that we have a biological system such as a population of water striders in Eldakar et al. in which intuitive inspection suggests that individual fitness depends on an individual component and a group component as in Figure 3 (a) and (b). Analysis reveals proposed causal pathways for individual trait and group trait to affect individual fitness via the two components. In particular, such an analysis comprises a hypothesis on the mechanism that mediates the effect of the group trait on the group component of individual fitness. For the water strider example the group trait is mean aggressiveness on a patch, the group component is proportional to the number of females on a patch, and the mechanism that mediates the effect of the former on the latter is female exodus determined by the females’ preference for low aggressiveness patches. Choosing contextual analysis or the Price approach for quantification goes hand in hand with the commitment to regard Figure 3 (a) or (b), resp., as the causal structure underlying the observed phenomena. The causal structures posited by the two approaches differ in that the Price approach assumes an effect of the group trait on the individual component of fitness. This assumption is reflected in the diagonal arrow in Figure 3 (b) that is missing in panel (a). The two arrows emanating from *X* in (b) represent two distinct cause-effect relations between the group trait and individual fitness. But given the hypothesis on the mechanism that mediates the effect of the group trait on the group component of fitness these two distinct cause-effect relations correspond to two distinct mechanisms through which the group trait affects fitness. Consequently, it is in principle possible to separate the effects by intervening on one of the mechanisms but not the other. This intervention translates to removing the vertical arrows from *X* to *w*_gr_ in Figure 3 (a) and (b) so that the system is described by the mutilated graphs in Figure 4. But in the system with suppressed group effects the two causal structures in Figure 4 (a) and (b) can be distinguished on the basis of the observable quantities *x*, *X*, and *w*. In particular, contextual analysis predicts individual fitness to be independent of group membership when the system is being intervened on in this way. The Price approach, however, predicts continued dependence of fitness on group trait due to within-group competition. As these predictions cannot both be true, the intervention allows the identification of one of the two approaches as being in accord with experimental observations.

**Figure 4:**
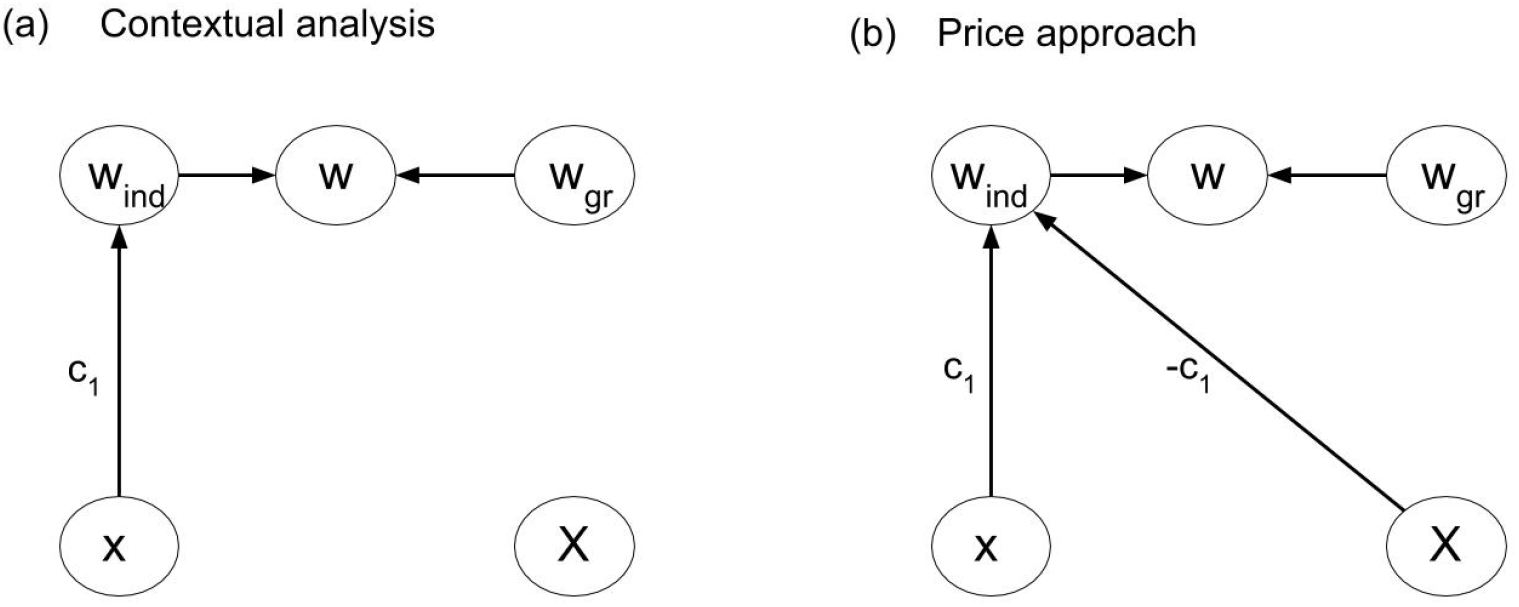
Mutilated causal graphs when suppressing the effect of the group trait on the group component. Contextual analysis predicts fitness to be independent of the group trait when the group effect is suppressed while the Price approach does not.

In the water strider example it is now easy to see how a decision for one of the two approaches may be reached. Since the effect of group trait on fitness is mediated by female exodus, the effect can be suppressed by preventing females from leaving patches, i.e., by removing female dispersal between patches (Eldakar et al., 2009). It is crucial that this intervention leaves the diagonal arrow in Figure 4 (b) intact. This is because the diagonal arrow represents a different causal pathway, namely the within-patch competition for females which is not affected by preventing females from leaving the patch. An informed decision for contextual analysis can then be reached if fitness is independent of mean aggressiveness on a patch when female dispersal is removed, i.e., if the diagonal arrow in Figure 4 (b) was not part of the underlying causal structure in the unperturbed system. The Price approach is more appropriate if fitness still depends on patch composition under this experimental condition.

Both the Price approach and contextual analysis serve the purpose to determine the quantities *w_ind_* and *w_gr_* in Equation (3), or equivalent quantities (see Table 1), from the more easily measurable variables individual trait and individual fitness. In order to achieve this, both approaches require assumptions that can be conveniently represented in terms of causal graphs as in Figure 3. We have shown above how, in principle, it is possible to determine which of the two approaches is more appropriate. However, we have seen that the causal structures posited are highly contrived. It seems therefore very well possible that neither of the two approaches is suitable for determining the level-specific strength of selection. This is the case when neither of the causal graphs (a) and (b) in Figure 3 represents the causal structure underlying the biological phenomenon in question.

## 4 Conclusion

Group selection refines kin selection by splitting individual fitness into two components, i.e., by assuming that fitness is determined by two additional factors that are themselves determined by the variables individual trait and group trait. The causal graphs in Figure 3 show that this means that group selection adds a layer to the causal structure of selection assumed by kin selection. This addition constitutes a proper refinement of kin selection and corresponds to avoiding averaging over the causes of individual fitness (the ‘averaging fallacy’ described by Sober and Wilson (1999)). From this viewpoint, the tension between contextual analysis and the Price approach can be seen as an instance of the purely formal problem of connecting an additional layer of nodes to an existing graph. The connection schemes proposed by contextual analysis and the Price approach, i.e., the coefficients of the paths targeting *w*_ind_ and *w*_gr_ in Figure 3, are two solutions to this problem. Since omitted paths in a causal graph represent hypotheses about the absence of effects the correct approach is the approach whose hypotheses are satisfied in the biological system at hand. Phrasing the problem in terms of causal graphs demonstrates that, even in the additive case, other refinements are in principle possible and could apply to scenarios in which the individual component is given neither by soft selection (Price approach) nor by hard selection (contextual analysis) but by intermediate selection regimes (Débarre and Gandon, 2011). Casting an MLS analysis in terms of refinements of causal graphs gives a formal argument for the non-equivalence of MLS and kin selection. We have argued that the refinement introduced by MLS is non-trivial (see difficulties with Price approach and contextual analysis) and provides a view on the system that is tailored to the levels of organisation in the system. This view is crucial when cause-effect relations that pertain to a specific level are manipulated or undergo change and the system-level consequences of such alterations are to be predicted. Strengthening the formal core of MLS not only facilitates the application of MLS in evolutionary science but also aids in assessing benefits, limitations, and formal requirements of this approach to empirical and theoretical biological scenarios.

## 5 Acknowledgements

We thank Miguel Brun-Usan, Jamie Caldwell, Laura Mears, Freddie Nash, Samir Okasha, Dave Prosser, and Michael Wade for helpful discussions and comments.

